# Early postnatal development of the cellular and circuit properties of striatal D1 and D2 spiny projection neurons

**DOI:** 10.1101/413740

**Authors:** Rohan N. Krajeski, Anežka Macey-Dare, Fran van Heusden, Farid Ebrahimjee, Tommas J. Ellender

## Abstract

A dysfunctional striatum is thought to contribute to neurodevelopmental disorders such as ADHD, Tourette’s syndrome and OCD. Insight into these disorders is reliant on an understanding of the normal development of the striatal cellular and circuit properties. Here we combined whole-cell patch-clamp electrophysiology and anatomical reconstructions of D1 and D2 striatal projection neurons (SPNs) in brain slices to characterize the development of the electrophysiological and morphological properties as well as their long-range and local inputs during the first three postnatal weeks. Overall, we find that many properties develop in parallel but we make several key observations. Firstly, that the electrophysiological properties of young D1 SPNs are more mature and that distinctions between D1 and D2 SPNs become apparent in the second postnatal week. Secondly, that dendrites and spines as well as excitatory inputs from cortex develop in parallel with cortical inputs exhibiting a prolonged period of maturation involving changes in postsynaptic glutamate receptors. Lastly, that initial local connections between striatal SPNs consist of gap junctions, which are gradually replaced by inhibitory synaptic connections. Interestingly, relative biases in inhibitory synaptic connectivity seen between SPNs in adulthood, such as a high connectivity between D2 SPNs, are already evident in the second postnatal week. Combined, these results provide an experimental framework for future investigations of striatal neurodevelopmental disorders and show that many of the cellular and circuit properties are established in the first and second postnatal weeks suggesting intrinsic programs guide their development.

**Significance Statement:** Normal brain development involves the formation of neurons, which develop correct electrical and morphological properties and are precisely connected with each other in a neural circuit. In neurodevelopmental disorders these processes go awry leading to behavioral and cognitive problems later in life. Here we provide for the first time a detailed quantitative description of the cellular and circuit properties of the two main neuron types of the striatum during the first postnatal weeks. This can form an experimental framework for future studies into neurodevelopmental disorders. We find that most of the properties for both types of striatal neuron develop in parallel and are already established by the second postnatal week suggesting a key role for intrinsic programs in guiding their development.

## Introduction

The striatum is the main input nucleus of the basal ganglia and consists of two populations of projection neurons with distinct long-range outputs: the D1-expressing direct pathway spiny projection neurons (SPNs) and the D2-expressing indirect pathway SPNs (Day *et al.*, 2008; Gertler *et al.*, 2008) that can differentially regulate motor behavior (Graybiel *et al.*, 1994; Grillner *et al.*, 2005; Yin & Knowlton, 2006; Kravitz *et al.*, 2010; Tecuapetla *et al.*, 2016). Adult D1 and D2 SPNs exhibit distinct electrical and morphological properties (Gertler *et al.*, 2008) and form precise non-random local synaptic connections with each other (Taverna *et al.*, 2008; Planert *et al.*, 2010; Cepeda *et al.*, 2013). Imbalances in the activity of the two pathways is thought to contribute to the cognitive and motor symptoms seen in late onset neurodegenerative disorders such as Parkinson’s disease (Taverna *et al.*, 2008) and Huntington’s disease (Cepeda *et al.*, 2013), but also those seen in early onset neurodevelopmental disorders such as Tourette’s syndrome (McNaught & Mink, 2011; Albin, 2018), OCD (Graybiel & Rauch, 2000; Langen *et al.*, 2011), ADHD (Del Campo *et al.*, 2011) and autism spectrum disorders (Shepherd, 2013). The cellular and neural circuit changes that underpin these neurodevelopmental disorders are major research areas. Although seminal papers have started to shed light on early postnatal striatal development (Tepper *et al.*, 1998; Dehorter *et al.*, 2011; Kozorovitskiy *et al.*, 2012; Peixoto *et al.*, 2016) many aspects of D1 and D2 SPN development have remained unstudied.

A combination of whole-cell patch-clamp electrophysiology and anatomical analysis in mouse brain slices allowed for the investigation of the cellular and circuit properties of striatal D1 and D2 SPNs at three key developmental time periods. These include postnatal day 3 to 6 (P3-6): this is the period when most striatal SPNs have been born but excitatory synaptic input to the striatum is thought to be minimal and mouse pups produce little movement, P9-12: when excitatory synaptic inputs to the striatum are thought to have undergone a period of rapid maturation and motor competence of the pups has increased and finally P21 and older (P21+): when the striatal neurons and the circuit are approaching maturity and mice readily traverse the environment (Tepper *et al.*, 1998; Khazipov *et al.*, 2004; Dehorter *et al.*, 2011; Kozorovitskiy *et al.*, 2012; Peixoto *et al.*, 2016). Firstly, we find that young D1 SPNs are electrophysiologically more mature than D2 SPNs and that intrinsic electrophysiological differences between D1 and D2 SPNs are already apparent during the second postnatal week. Secondly, we find that the dendrites and spines of D1 and D2 SPNs develop in parallel during the first postnatal weeks. Thirdly, we find that excitatory cortical synapses are functional in the first postnatal week and are equally sampled by D1 and D2 SPNs. Further maturation of excitatory inputs occurs in parallel including a progressive increase in AMPA receptor-mediated currents. Finally, simultaneous quadruple patch-clamp recordings allowed for the investigation of local connections between developing SPNs. This reveals that in the first postnatal week SPNs form gap junctions with each other unbiased to SPN type and that D1 SPNs are the first to form inhibitory synapses with other SPNs. In the second postnatal week gap junction incidence has halved and all SPNs are increasingly forming inhibitory synaptic connections with each other. Interestingly, the synaptic connections are precise and non-random with the majority of synaptic connections formed between D2 SPNs. Indeed, the relative biases in synaptic connectivity found in adulthood are already apparent in the second postnatal week. Together, these results suggest that many of the striatal cellular and circuit properties are already established before mice receive sensory input (Tobach *et al.*, 1971) or are freely exploring the environment (Dehorter *et al.*, 2011) suggesting a major role for intrinsic developmental programs.

## Results

### Development of the electrophysiological properties of D1 and D2 SPNs

We first investigated the development of intrinsic electrophysiological properties of striatal D1 and D2 SPNs and their ability to initiate action potentials using whole-cell current-clamp recordings of SPNs in mouse brain slices at postnatal (P) day 3-6, P9-12 and P21 and older (**Fig. 1A**). Mice consisted of heterozygous D1 and D2-GFP mice and wildtype C57/BL6 mice (see **Methods**) and we therefore always included biocytin in the intracellular solution, followed by immunocytochemistry for CTIP2 and PPE (Gerfen *et al.*, 1990; Arlotta *et al.*, 2008) to confirm, whenever possible, whether we recorded from D1 or D2 SPNs (**Fig. 1B**). Hyperpolarizing and depolarizing current steps were injected into the recorded SPNs to characterize their electrophysiological properties (**Fig. 1C** and **Table 1**).

**Table 1:**
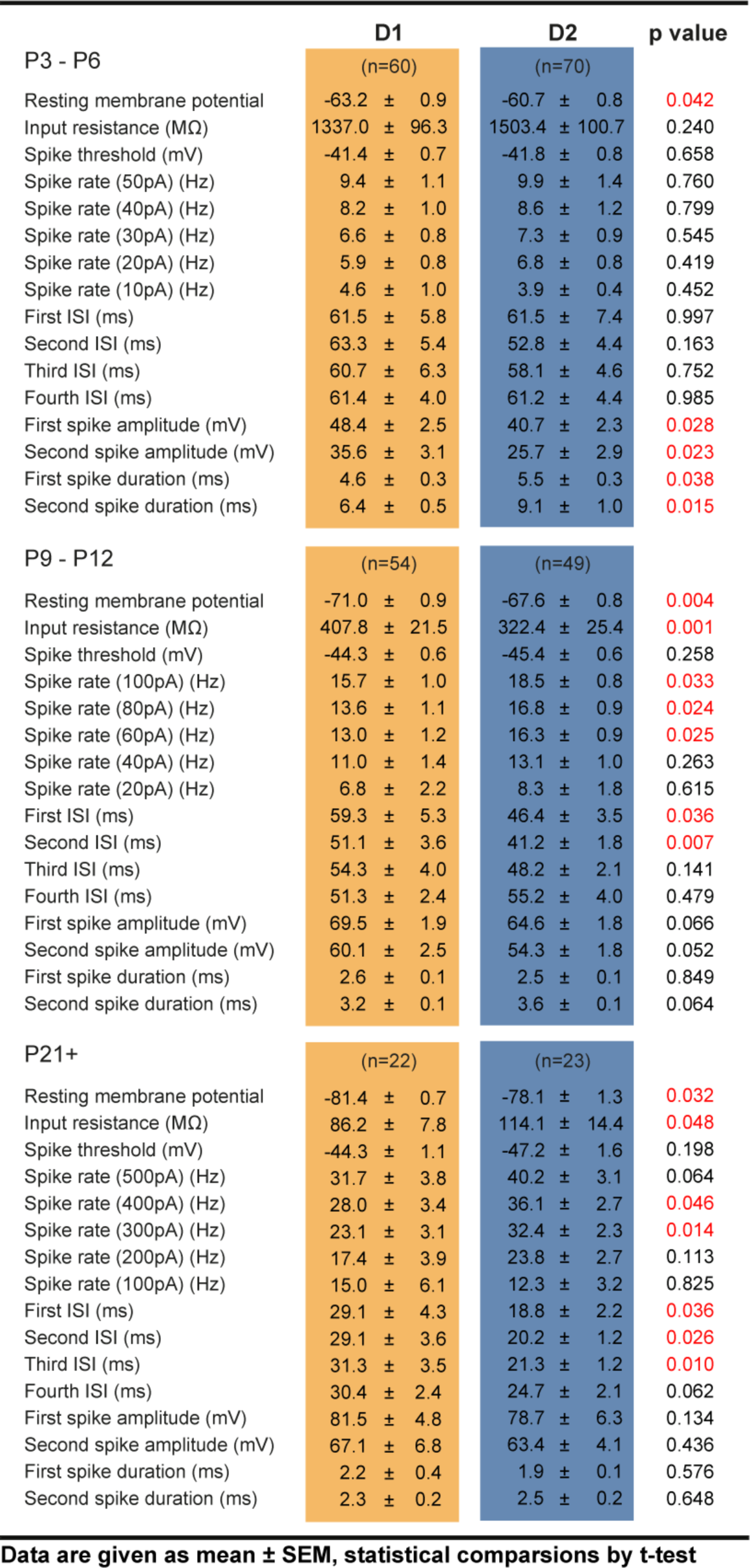
Intrinsic membrane properties of D1 and D2 SPNs

Firstly, we find that in the earliest postnatal period (P3-6) most SPNs (83.0%) are already able to initiate small amplitude action potentials (**Fig. 1C**). We find that the D1 SPNs exhibit subtle but significant maturational differences consistent with their suggested earlier birthdate (Marchand & Lajoie, 1986; van der Kooy & Fishell, 1987; Kelly *et al.*, 2018), including relatively larger and narrower action potentials (D1 amplitude first spike: 48.4 ± 2.5 mV and D2 amplitude first spike: 40.7 ± 2.3 mV, D1 amplitude second spike: 35.6 ± 3.1 mV and D2 amplitude second spike: 25.7 ± 2.9 mV, D1 first spike duration: 4.6 ± 0.3 ms and D2 first spike duration: 5.5 ± 0.3 ms, D1 second spike duration: 6.4 ± 0.5 ms and D2 second spike duration: 9.1 ± 1.0 ms, p=0.028, p=0.023, p=0.038 and p=0.015, t-test, D1 n=60 and D2 n=49, **Fig. 1D** and **Table 1**). These differences are transient and in later stages all SPNs generate large amplitude and narrow action potentials (**Table 1**). Secondly, we find that the passive membrane properties gradually change and include the emergence of a pronounced inward rectifying current at later developmental stages (**Fig. 1E**). Thirdly, we find we can elicit higher frequencies of action potentials as the SPNs mature using a range of depolarizing current steps chosen to allow sufficient depolarization of SPNs beyond spike threshold taking in consideration developmental changes in input resistance (D1; P3-6: 8.2 ± 1.0 Hz, P9-12 13.6 ± 1.1 Hz and P21+: 28.0 ± 3.4 Hz; P3-6 vs P9-12: p=0.00043 and P9-12 vs P21+: p=0.00013, Mann-Whitney U test, and D2; P3-6: 8.6 ± 1.2 Hz, P9-12 16.8 ± 0.9 Hz and P21+: 36.1 ± 3.7 Hz; P3-6 vs P9-12: p=6.23E-7 and P9-12 vs P21+: p=2.85E-8, Mann-Whitney U test, **Fig. 1F** and **G**, **Table 1**).

**Figure 1:**
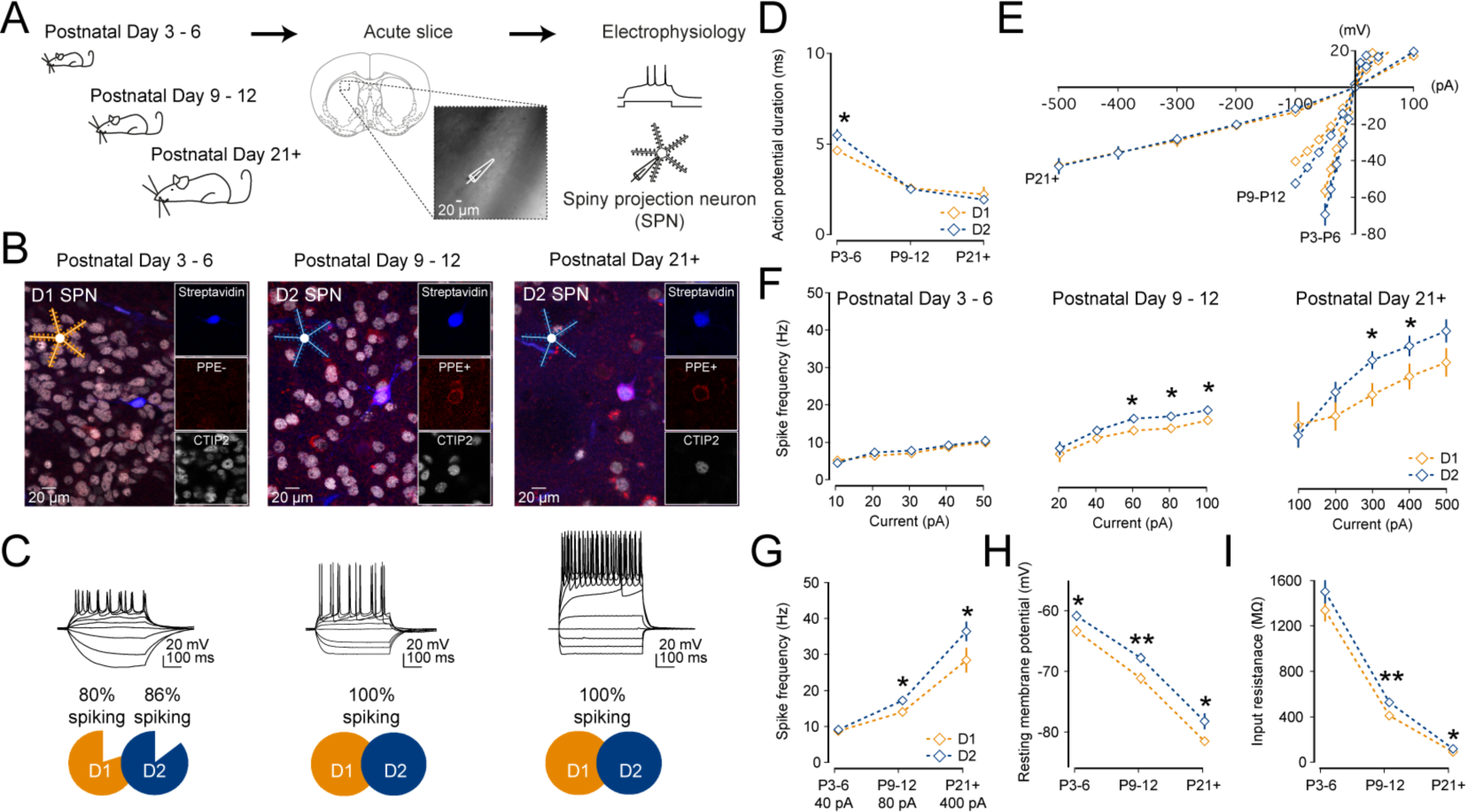
Maturational and intrinsic differences in electrophysiological properties of D1 and D2 SPNs. (**A**) Whole-cell patch-clamp recordings were made from spiny projection neurons (SPNs) in acute coronal brain slices of mice at three different time periods; Postnatal (P)3-6, P9-12 and P21 and older. (**B**) Internal recording solutions included biocytin allowing for posthoc confirmation of SPN type using immunocytochemistry for the SPN marker CTIP2 (chicken ovalbumin upstream promoter transcription factor-interacting protein-2) and D2 SPN marker PPE (pre-proenkephalin). Note the example P3-6 SPN is PPE-negative and CTIP2-positive, corresponding to a putative D1 SPN (in orange), whereas the example SPNs for both P9-12 and P21+ are PPE-positive and CTIP2-positive, corresponding to putative D2 SPNs (in blue). (**C**) Hyperpolarizing and depolarizing current steps were used to characterize the electrophysiological properties of SPNs. We find that the majority of P3-6 SPNs are able to generate small action potentials. (**D**) Action potential duration is significantly shorter for D1 SPNs at P3-6. (**E**) As the SPNs mature, a pronounced inward rectifying current develops. (**F**, **G**) Whereas in the first postnatal week both D1 and D2 SPNs exhibit a similar action potential frequency, at later stages in development the D2 SPNs start exhibiting a significantly higher action potential frequency, which persists into adulthood. (**H**) The resting membrane potential becomes progressively more hyperpolarized as SPNs mature. Note the consistently more depolarized potential of the D2 SPNs relative to the D1 SPNs. (**I**) The input resistance decreases as SPNs mature. Note the consistently higher input resistance of the D2 SPNs relative to the D1 SPNs.

We find that the D2 SPNs consistently exhibit a higher firing frequency at later developmental stages (P9-12 at 80 pA; D1: 13.6 ± 1.1 Hz and D2: 16.8 ± 0.9 Hz, P21+ at 400 pA; D1: 28.0 ± 3.4 Hz and D2: 36.1 ± 2.7 Hz, p=0.024 and p=0.046, t-test, **Fig. 1F** and **G**, **Table 1**). The higher action potential frequency seen during later developmental stages is likely aided by the consistently more depolarized membrane potential (P9-12; D1: −71.0 ± 0.9 mV and D2: −67.6 ± 0.8 mV, p=0.004 and p21+; D1: −81.4 ± 0.7 mV and D2: 78.1 ± 1.3 mV, p=0.032, t-test, **Fig. 1H**) and their higher input resistance (P9-12; D1: 407.8 ± 21.5 MΩ and D2: 322.4 ± 25.4 MΩ, p=0.001 and P21+; D1: 86.2 ± 7.8 MΩ and D2: 114.1 ± 14.4 MΩ, p=0.048, t-test, **Fig. 1I**) of the D2 SPNs.

In conclusion, we find that most D1 and D2 SPNs are able to generate action potentials shortly after birth. Maturational differences can be seen early on including narrower and larger action potentials in the D1 SPNs, which disappear soon after. Furthermore, significant differences in the intrinsic membrane properties can already be observed in the second postnatal week, which persist into adulthood and include a higher excitability and action potential frequency of the D2 SPNs.

### Development of dendritic morphological properties of D1 and D2 SPNs

We next investigated the development of the dendritic morphology of SPNs allowing them to receive and process excitatory and inhibitory inputs. The addition of biocytin in internal recording solutions allowed for subsequent DAB immunohistochemistry and reconstruction of dendritic arbors of previously recorded neurons (**Fig. 2A**). We find that the dendritic length of both D1 and D2 SPNs more than doubles (from ∼700 μm to ∼1800 μm) as the neurons mature (D1; P3-6: 698.4 ± 97.6 μm, P9-12: 1556.1 ± 181.6 μm and P21+: 1907.63 ± 166.5 μm, p=0.00022 and p=0.164, Mann-Whitney U test, n=21, 10 and 19 and D2; P3-6: 757.4 ± 119.0 μm, P9-12: 1350.1± 83.9 μm and P21+: 2109.7 ± 124.3 μm, p=0.00088 and p=0.00031, Mann-Whitney U test, n=22, 13 and 28; **Fig. 2B**), but we find no significant difference in the length of the dendritic arbor between the D1 and D2 SPNs at the various ages (P3-6: p=0.827, P9-12: p=0.738 and P21+: p=0.269; Mann-Whitney U test). This increase in dendritic arbor was concomitant with a significant increase in distal dendritic complexity allowing the SPNs to sample larger and extensive regions of striatum (P3-6 vs P9-12: F(17, 691)=8.98, p=2.85E-21 and P9-P21 vs P21+: F(22,1078)=4.136, p=7.02E-10, ANOVA) but similarly no significant differences were observed between the D1 and D2 SPNs at the various ages (P3-6: p=0.816, P9-12: p=0.091 and P21+: p=0.235, ANOVA; **Fig. 2C**). The orientation of the dendritic branches of both D1 and D2 SPNs exhibited a predominantly radial morphology from birth onwards but we observed a preferred lateral-ventral to a medial-dorsal orientation (**Fig. 2A** and **D**).

**Figure 2:**
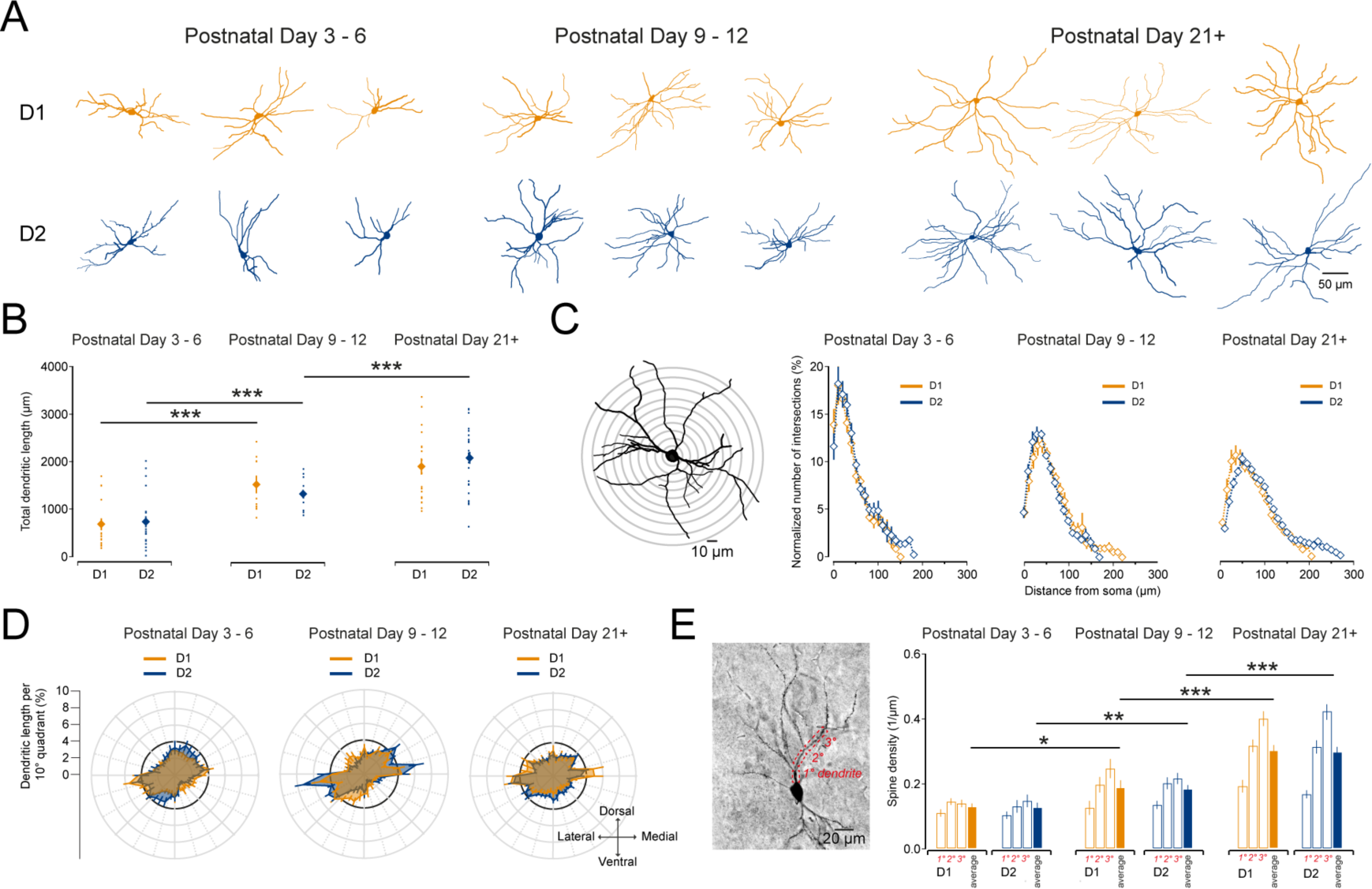
Development of dendritic arbors and spines of D1 and D2 SPNs. (**A**) Example reconstructions of previously recorded SPNs which were processed for DAB immunohistochemistry. SPNs are grouped according to age (left to right; P3-6, P9-12 and P21+) and whether they are D1 (orange, top) or D2 SPNs (blue, bottom). The examples shown and all reconstructed and analyzed neurons are from coronal sections and are aligned such that top is dorsal, bottom is ventral, left is lateral and right is medial. (**B**) D1 (orange) and D2 SPNs (blue) exhibit a significant and similar increase in their dendritic length as they mature (**C**) Scholl analysis of dendritic complexity of D1 and D2 SPNs reveals a similar elaboration of distal dendritic segments as they mature. (**D**) Polarity analysis of dendrites of D1 and D2 SPNs reveal a mostly uniform and radial distribution of their dendrites. Note the bias to extend dendrites from lateral-ventral aspects to medial-dorsal aspects. (**E**) Spine density of D1 and D2 SPNs at different ages. Note the similar increase in spine density in both D1 and D2 SPNs as they mature.

Lastly, we investigated the density of dendritic spines on primary, secondary and tertiary dendritic branches on both D1 and D2 SPNs at the three ages (**Fig. 2E**). Whereas we see a significant increase in the average spine density (per μm of dendrite) as the neurons mature (D1; P3-6: 0.13 ± 0.01, P9-12: 0.19 ± 0.02 and P21+: 0.30 ± 0.02, p=0.02 and p=0.0006, Mann-Whitney U test, n=8, 8 and 10 and D2; P3-6: 0.13 ± 0.02, P9-12: 0.18 ± 0.01 and P21+: 0.30 ± 0.02, p=0.008 and p=0.0002, Mann-Whitney U test, n=10, 11 and 18), we did not see a significant difference between the D1 and D2 neurons at the various ages (p>0.05).

In conclusion, we find that the development of the morphology of the D1 and D2 SPNs run in parallel with no observed differences in the development of their dendritic arbors or spine density.

### Maturation of excitatory and inhibitory synaptic inputs onto SPNs

Our results so far suggest that the D1 and D2 SPNs are already able to initiate action potentials during the first postnatal week and that the development of their dendritic arbors and spines develops mostly in parallel allowing them to sample excitatory and inhibitory inputs from nearby axons. We next characterized the development of the functional synaptic inputs to the SPNs by performing whole-cell voltage-clamp recordings of SPNs in the presence of TTX (1 μM). This allowed us to record spontaneous miniature excitatory postsynaptic currents (mEPSCs) by holding the SPN membrane voltage at −70 mV (**Fig. 3A**) and spontaneous miniature inhibitory postsynaptic currents (mIPSCs) by holding the SPN membrane voltage at 0 mV (**Fig. 3D**). Our first observation is that excitatory inputs as mEPSCs can be detected as early as P3-6 in both D1 and D2 SPNs (**Fig. 3A**, **B**) and although there is a trend towards an increase across developmental time (P3-6: 0.85 ± 0.10 Hz and p21+: 1.01 ± 0.09 Hz, p=0.319, Mann-Whitney U test, n=15 and n=22, **Fig. 3B**) their frequency is already close to that seen in adulthood (∼1 Hz). In contrast, we find that the mEPSC amplitude significantly increases for both D1 and D2 SPNs from P3-6 to P9-12 (D1; P3-6: 3.40 ± 0.40 pA to D1: P9-12: 8.38 ± 0.64 pA and D2; P3-6: 3.29 ± 0.74 pA to D2: P9-12: 7.68 ± 0.49 pA; p=0.005 and p=0.003, Mann-Whitney U test, both n=5 and n=7; **Fig. 3C**) after which it stays constant. We did not find significant differences in either the mEPSC frequency or amplitude between the D1 and D2 SPNs at any of the age ranges investigated (p>0.05). These results suggest that excitatory inputs are present and functional soon after birth and develop in parallel and similarly innervate both D1 and D2 SPNs with pronounced postsynaptic changes occurring between P3-6 and P9-12.

The response of SPNs to excitatory input is modulated by concurrent inhibitory inputs that they might receive. We next investigated the development of inhibitory synaptic inputs onto SPNs as reflected in the frequency and amplitude of mIPSCs. We find that both D1 and D2 SPN receive inhibitory synaptic input starting from P3-6 onwards (**Fig. 3D**, **E**). However, the frequency of the mIPSCs at this age range is comparatively low at ∼0.1 Hz and exhibits a steady increase across all three age ranges (D1; P3-6: 0.10 ± 0.02 Hz, P9-12: 0.35 ± 0.05 Hz and P21+: 0.68 ± 0.08 Hz; p=0.005 and p=0.013, Mann-Whitney U test, n=5, 7 and 8 and D2; P3-6: 0.14 ± 0.02 Hz, P9-12: 0.30 ± 0.04 Hz and P21+: 0.63 ± 0.15 Hz; p=0.018 and p=0.048, Mann-Whitney U test, n=5, 7 and 6; **Fig. 3E**).

**Figure 3:**
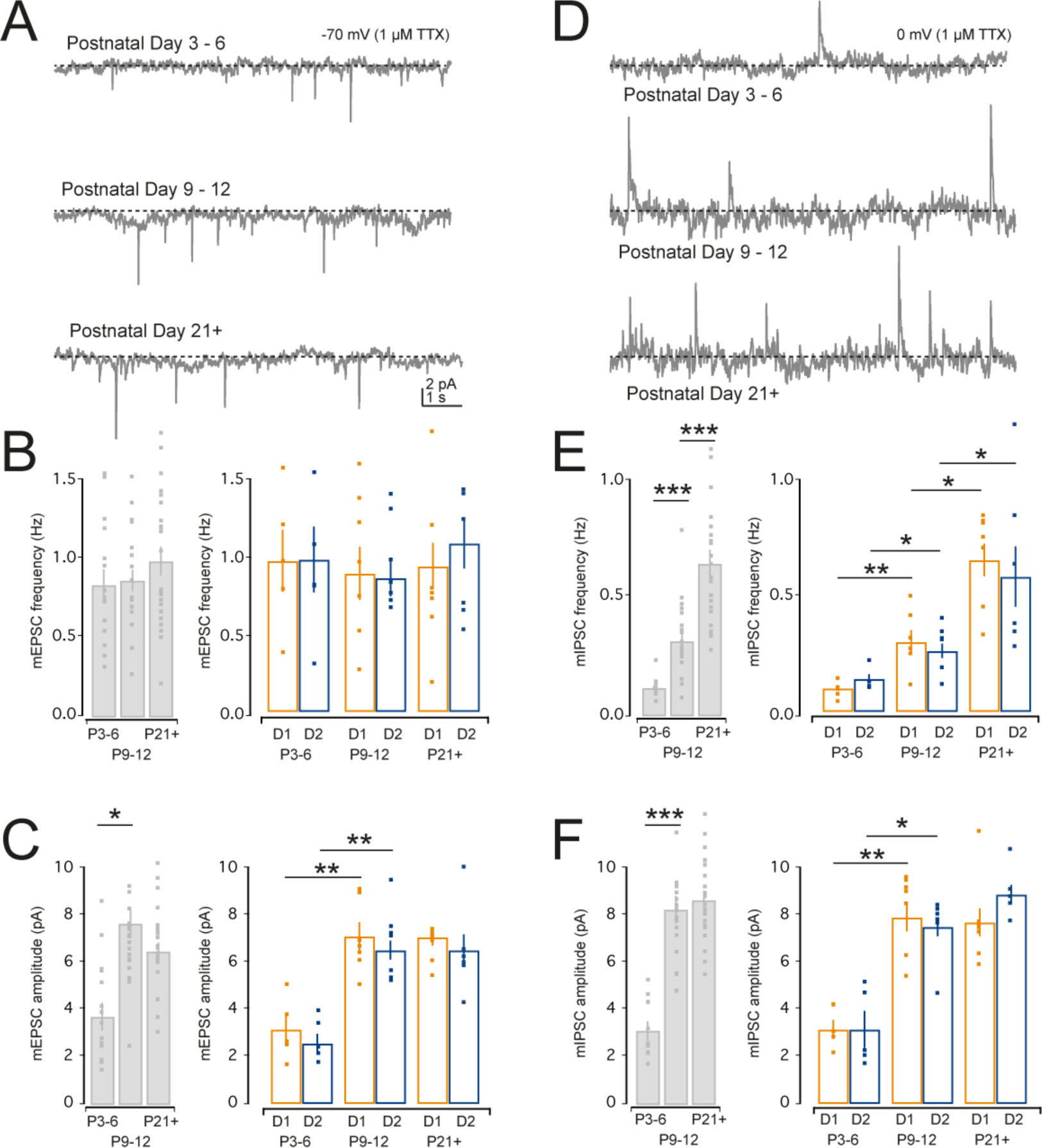
Characterization of miniature EPSCs and IPSCs in D1 and D2 SPNs. (**A**) Miniature EPSCs (mEPSCs) were recorded as downward deflections from SPNs held in voltage-clamp at a holding potential of −70 mV in the presence of TTX (1 μM). (**B**) Bar plot of mEPSC frequency showing a relatively stable mEPSC frequency across developmental time for all recorded SPNs (left) and identified SPNs (right). Note the absence of significant differences in mEPSC frequency between D1 (orange) and D2 SPNs (blue). (**C**) Bar plot of mEPSC amplitude for all recorded SPNs showing a significant increase in amplitude between P3-6 and P9-12 (p=0.017, left). This increase in amplitude is seen for both the D1 and D2 SPNs (right). (**D**) Miniature IPSCs (mIPSCs) were recorded as upward deflections from SPNs held in voltage-clamp at a holding potential of 0 mV in the presence of TTX (1 μM). (**E**) Bar plot of mIPSC frequency shows a steady increase in mIPSC frequency across developmental time (P3-P6 to P9-P12, p=0.00002 and P9-P12 to P21+, p=0.000014, left) which is seen for both D1 and D2 SPNs (right). (**F**) Bar plot of mIPSC amplitude showing a significant increase in mIPSC amplitude between P3-6 and P9-12 (p=2.16E-8, left) for both D1 and D2 SPNs (right).

Similar to the mEPSC amplitude, we find that the mIPSC amplitude also exhibits a significant increase from P3-6 to P9-12 (D1; P3-6: 3.19 ± 0.60 pA and P9-12: 7.17 ± 0.57 pA and D2; P3-6: 2.79 ± 0.42 pA and P9-12: 6.93 ± 0.59 pA; p=0.003 and p=0.01, Mann-Whitney U test, both n=5 and n=7; **Fig. 3F**) after which it does not further increase.

These results suggest that functional excitatory and inhibitory synaptic inputs are present during the first postnatal week and similarly develop for both the D1 and D2 SPNs. Moreover, they suggest that between the first and second postnatal weeks substantial postsynaptic changes occur as reflected in the increased mEPSC and mIPSC amplitudes. Whereas the frequency of mEPSCs stays relatively constant, we find that the mIPSC frequency increases over time suggesting a prolonged presynaptic development of inhibitory inputs.

### Maturation of long-range cortical excitatory synapses on striatal SPNs

The main excitatory afferents to striatal SPNs arise from neurons located in the cortex and thalamus (Kemp & Powell, 1971; Buchwald *et al.*, 1973; Smith *et al.*, 2004) and anatomical work suggests their axons and synapses are present in the striatum early on in development (Nakamura *et al.*, 2005; Sohur *et al.*, 2012) but when and to what extent they are sampled by D1 and D2 SPNs is unknown. We performed whole-cell patch-clamp recordings of identified SPNs in the dorsal striatum and activated excitatory afferents from cortex by giving single and trains of stimulation via a Tungsten bipolar stimulating electrode placed in the external capsule (**Fig. 4A**).

Firstly, we find that not all SPNs at P3-6 receive cortical excitatory synaptic inputs whereas those at P9-12 and older all did (P3-6; D1: 75% and D2: 71%, n=9 and n=10, **Fig. 4B**). For all the SPNs that do receive an input we find that across a wide range of stimulation strengths (range 20-220 μA) both the D1 and D2 SPNs receive similar amplitude EPSPs at all three age ranges (P3-6: *F* (1, 83) = 0.012, p=0.918, P9-12: *F* (1, 94) = 0.262, p=0.610 and P21+: *F* (1, 51) = 1.414, p=0.240). Similarly, comparing the maximum EPSP amplitudes that we could elicit in SPNs across the age ranges shows relatively stable EPSP amplitudes (P3-6: 2.86 ± 0.68 mV, P9-12: 3.51 ± 0.83 mV and P21+: 3.54 ± 0.73 mV, P3-P6 vs P21+, p=0.129, Mann-Whitney U test, n=19, 20 and 17, **Fig. 4C**). However, when we record in voltage-clamp mode (at −75 mV) we find a robust increase in elicited EPSC amplitude (P3-6: 24.10 ± 6.61 pA, P9-12: 31.18 ± 11.87 pA and P21+: 73.42 ± 12.01 pA, P3-6 vs P21+, p=0.001 and P9-12 vs P21+, p=0.017, Mann Whitney test, n=14, 10 and 29, **Fig. 4C**) suggestive of a progressive strengthening of corticostriatal synapses.

**Figure 4:**
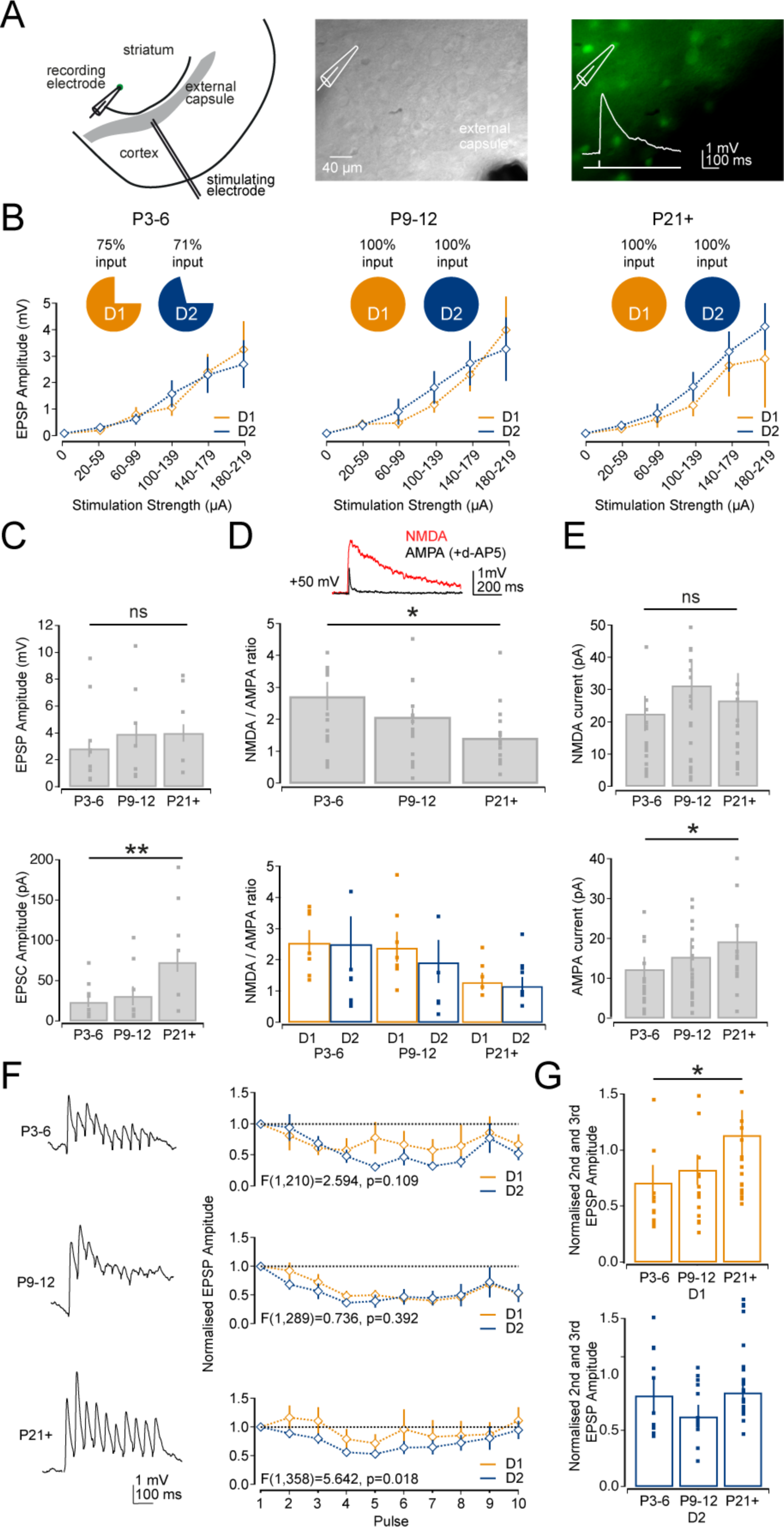
Development of excitatory cortical synaptic inputs onto D1 and D2 SPNs. (**A**) Diagram of recording configuration and placement of stimulation electrode to activate cortical afferents (left). Example Dodt contrast image (middle) and fluorescence image (right) of striatum in a D2-GFP transgenic mouse. Inset: example of an evoked cortical EPSP. (**B**) Graphs of EPSP amplitude across a range of stimulation strengths (range 20-220 μA) for the three different age ranges. Note the similar amplitude in evoked EPSPs between the D1 (orange) and D2 SPNs (blue). Note also that ∼70% of SPNs exhibited a response at P3-6 whereas they always exhibited a response at later age ranges. (**C**) Bar plot of the maximum evoked EPSP amplitude across the three age ranges which remains relatively constant at ∼3 to 4 mV (top). Bar plot of EPSC amplitude shows an increase in the cortically evoked excitatory current (bottom). (**D**) Bar plot of the NMDA/AMPA receptor ratio across the age ranges. Note the significant decrease in the ratio as the neurons mature (top), which occurs in parallel for both D1 and D2 SPNs (bottom). (**E**) Bar plot of the NMDA receptor-mediated current (top) and AMPA receptor-mediated current (bottom) suggesting the change in NMDA/AMPA ratio is a reflection of a significant increase in the AMPA receptor-mediated current. (**F**) Graphs of the EPSP amplitude across 10 stimulations at 20Hz showing that corticostriatal synapses at D1 and D2 SPNs predominantly exhibit short-term depression at all three age ranges with corticostriatal synapses on D2 SPNs exhibiting more pronounced depression than those onto D1 SPNs at P21+. (**G**) Barplots of the combined second and third normalized EPSP amplitude for the D1 (top) and D2 (bottom) SPNs. Note that the short-term plasticity at corticostriatal synapses onto D1 SPNs progressively change to neutral / facilitating across development which is not observed for the D2 SPNs.

Indeed, an increase in amplitude of the evoked excitatory response could be the result of changes in postsynaptic glutamate receptors. We next investigated the contribution of both NMDA and AMPA glutamate receptors to the cortically evoked excitatory responses at these different age ranges. Evoked excitatory events were recorded from SPNs in whole-cell voltage-clamp mode at a holding potential of +50 mV and consisted of combined NMDA and AMPA receptor-mediated currents (**Fig. 4D**). After baseline recording, d-AP5 (50 μm) was bath applied to block the NMDA receptor-mediated currents, thereby isolating the AMPA receptor-mediated component. Analysis of the ratio of peak amplitude NMDA and AMPA receptor-mediated currents showed a gradual and progressive decline in the NMDA/AMPA receptor ratio across developmental time (P3-6: 2.73 ± 0.45, P9-12: 2.08 ± 0.25 and P21: 1.43 ± 0.21, P3-6 vs P21+: p=0.046, Mann-Whitney U test, n=21, n=23 and n=17, **Fig. 4D**) similarly for both the D1 and D2 SPNs (**Fig. 4D** and **Table 2**). This change in the NMDA/AMPA receptor ratio is reflected in a progressive increase in AMPA receptor-mediated current (P3-6: 12.37 ± 3.05 pA, P9-12: 15.45 ± 3.77 pA and P21+: 19.31 ± 3.41 pA, P3-6 vs P21+: p=0.039, Mann-Whitney U test, n=21, n=23 and n=17) whereas the NMDA receptor-mediated current does not change significantly (P3-P6: 22.62 ± 5.51 pA, P9-12: 31.35 ± 7.69 pA and P21+: 26.72 ± 8.41 pA, P3-6 vs P9-12: p=0.562, P3-P6 vs P21+: 0.750, Mann-Whitney U test, n=21, n=23 and n=17, **Fig. 4E**).

**Table 2:**
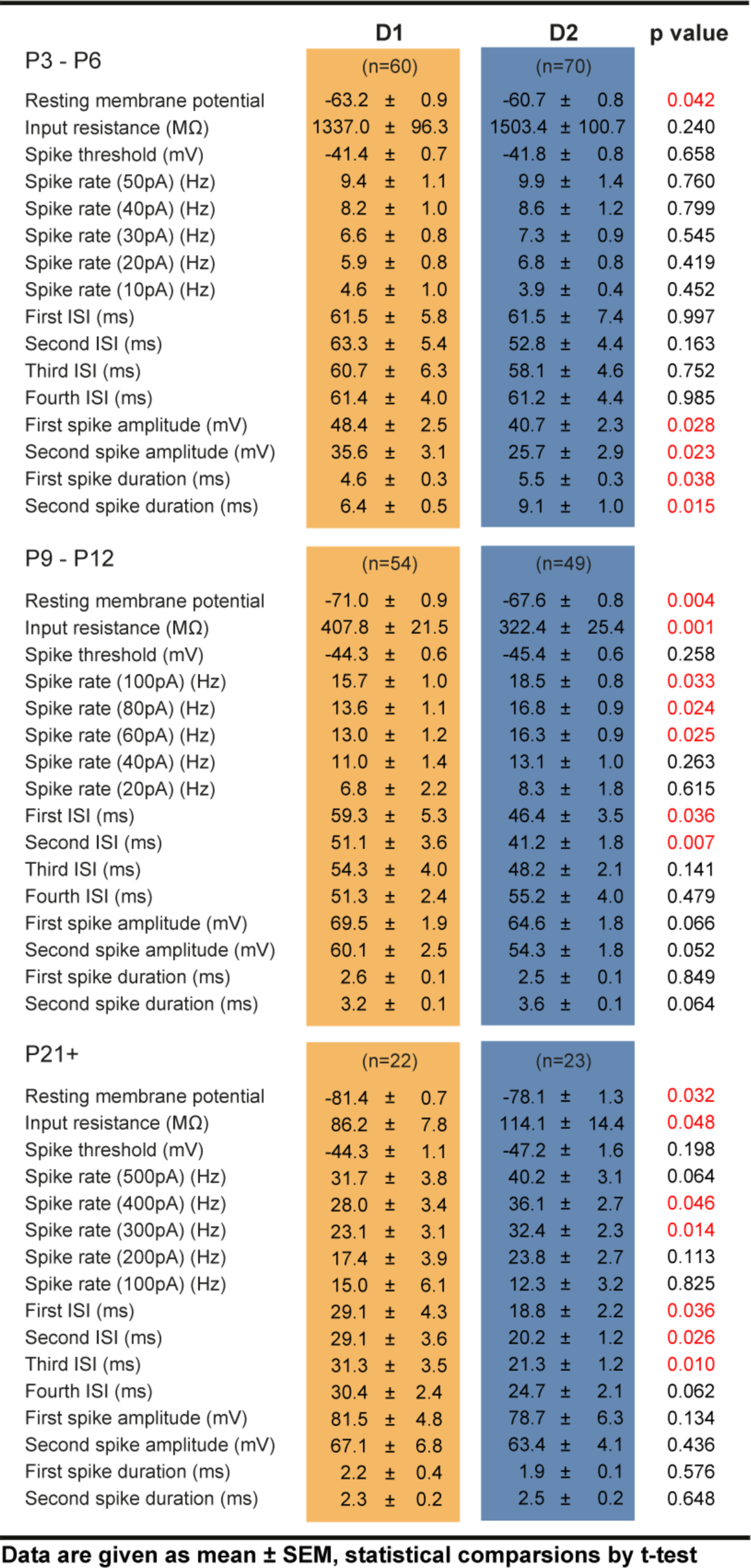
Properties of excitatory cortical synapses on D1 and D2 SPNs

Changes in postsynaptic ionotropic glutamate receptors are predicted to change the EPSP kinetics (Seeburg, 1993). Indeed, we find a faster rise time, decay time and duration of EPSPs at P21+ as compared to P3-6 (see **Table 2**). Interestingly, we find a transient phase at P9-12 where the EPSP duration is markedly longer than at the other ages (All SPNs; P3-6: 164.07 ± 21.25 ms, P9-12: 201.82 ± 12.53 ms and P21+: 118.42 ± 11.92 ms; P3-6 vs P9-12: p=0.046 and P9-12 vs P21+: p=0.00012, Mann-Whitney U test n=19, 20 and 17) as a result of a longer decay time of the EPSP (All SPNs; P3-6: 81.14 ± 9.66 ms, P9-12: 103.55 ± 7.29 ms and P21+: 58.72 ± 5.92 ms; P3-6 vs P9-12: p=0.046 and P9-12 vs P21+: p=0.00015, Mann-Whitney U test n=19, 20 and 17, **Table 2**) consistent with previous observations suggesting a transient expression of NR2C/D subunit containing NMDA receptors with longer kinetics (Monyer *et al.*, 1994; Dehorter *et al.*, 2011).

Lastly, we investigated the short-term plastic properties of the cortical synapses on SPNs using trains of electrical stimulation (10 pulses at 20 Hz). We find that cortical synapses are predominantly depressing at all developmental ages and are more depressing for D2 SPNs at P21+ (P3-6: *F* (1, 210) = 2.594, p=0.109, P9-12: *F* (1, 289) = 0.736, p=0.392 and P21+: *F* (1, 358) = 5.642, p=0.018, **Fig. 4F**). Indeed, when we compare the short-term plasticity of the combined second and third normalized EPSP amplitude across the age ranges we observe a progressive change from depressing towards neutral or facilitating for D1 SPNs whereas synapses onto D2 SPNs remain depressing (**Fig. 4G**).

Combined, these results suggest that the excitatory cortical inputs to D1 and D2 SPNs are functional in the first postnatal week and develop in parallel including a progressive strengthening of corticostriatal synapses through an increase in AMPA receptor-mediated transmission.

### Maturation of local inhibitory synaptic connections between striatal SPNs

Our data so far suggests that the majority of D1 and D2 SPNs sample excitatory input from cortex and are able to generate action potentials during the first postnatal week. However, the likelihood and timing of action potential generation is under control of local inhibition provided by both lateral inhibitory connections between SPNs as well as input from striatal interneurons (Tepper & Plenz, 2006; Ponzi & Wickens, 2010). Our analysis of mIPSC frequency would suggest an extended and progressive strengthening of inhibitory inputs across the age ranges. To investigate when SPNs form inhibitory synaptic connections with each other and how these change over development we performed quadruple whole-cell current-clamp recordings of SPNs at the three developmental age ranges (**Fig. 5A**) including immunocytochemistry (**Fig. 5B**) and histochemistry (**Fig. 5C**) as previously described to classify recorded neurons as D1 or D2 SPNs (see **Methods**). As immature SPNs also form gap junctions with each other (Venance *et al.*, 2004) we used hyperpolarizing and depolarizing current steps to investigate both gap junctions (**Fig. 5D**) as well synaptic connections (**Fig. 5E**) between SPNs.

We find that young SPNs form gap junctions with each other early in development (P3-6: 3.5%, n=13/377) but seem to progressively lose these connections and we were unable to detect gap junctions at P21+ (P9-12: 1.9% and P21+: 0%, 8/418 and 0/134, P3-6 vs P21+ p=0.0462, Fisher exact test, **Fig. 5F**). The majority of detected gap junctions were symmetrical (P3-6: 76.92% and P9-12: 75%) consistent with previous observations (Venance *et al.*, 2004). When we split the data into the various SPN connection groups; D1 to D1, D1 to D2, D2 to D1 and D2 to D2, we find that gap junction incidence does not differ between the various groups (**Fig. 5G**) but find that consistently the highest gap junction incidence can be found between D2 SPNs (P3-6: 3.45% and P9-12: 1.96%, **Fig. 5G**). We did not find significant differences in either the amplitude, rise time, coupling coefficient or junction conductance of gap junctions between the different SPN connection groups (**Table 3**). Although we observed an increase in the junction conductance from P3-6 to P9-12 (P3-6: 327.90 ± 64.56 and P9-12: 1911.85, p=0.00017, Mann-Whitney U test) the overall strength of the gap junctions and the impact of junctional currents seemed to decrease over time as reflected by a trend towards a decreased coupling coefficient (P3-6: 3.04 ± 0.61 and P9-12: 1.55 ± 0.32, p=0.072, Mann-Whitney U test, n=11 and n=10, **Table 3**) as has been reported for other brain regions (Yu *et al.*, 2012; Belousov & Fontes, 2013) and most likely the result of development changes in SPN resistance and capacitance.

**Table 3:** Properties of gap junctions between D1 and D2 SPNs

|  | D1<br>to<br>D1 | D1<br>to<br>D2 | D2<br>to<br>D2 | D2<br>to<br>D1 | All |
| --- | --- | --- | --- | --- | --- |
| P3 - P6 |  |  |  |  |  |
| Amplitude (mV) | 1.25 ± 0.18 | 1.02 ± 0.22 | 1.19 ± N/A | 0.86 ± N/A | 0.92 ± 0.13 |
| Rise time (ms) | 116.45 ± 14.65 | 125.75 ± 2.75 | 125.00 ± N/A | 146.50 ± N/A | 107.02 ± 8.57 |
| Coupling coefficient (%) | 1.50 ± 0.07 | 6.28 ± 1.31 | 4.40 ± N/A | 3.03 ± N/A | 3.04 ± 0.61 |
| Junctional conductance (pS) | 194.02 ± 12.96 | 221.82 ± 52.56 | 182.88 ± N/A | 605.25 ± N/A | 327.90 ± 64.56 |
| P9 - P12 |  |  |  |  |  |
| Amplitude (mV) | N/A ± N/A | 2.37 ± N/A | 0.48 ± 0.21 | 2.16 ± N/A | 1.11 ± 0.30 |
| Rise time (ms) | N/A ± N/A | 83.60 ± N/A | 97.10 ± 17.80 | 85.20 ± N/A | 106.88 ± 13.66 |
| Coupling coefficient (%) | N/A ± N/A | 3.33 ± N/A | 1.51 ± 0.71 | 2.64 ± N/A | 1.55 ± 0.32 |
| Junctional conductance (pS) | N/A ± N/A | 385.25 ± N/A | 2612.83 ± 1146.57 | 411.29 ± N/A | 1911.85 ± 407.34 |
| P21+ |  |  |  |  |  |
| Amplitude (mV) | N/A ± N/A | N/A ± N/A | N/A ± N/A | N/A ± N/A | N/A ± N/A |
| Rise time (ms) | N/A ± N/A | N/A ± N/A | N/A ± N/A | N/A ± N/A | N/A ± N/A |
| Coupling coefficient (%) | N/A ± N/A | N/A ± N/A | N/A ± N/A | N/A ± N/A | N/A ± N/A |
| Junctional conductance (pS) | N/A ± N/A | N/A ± N/A | N/A ± N/A | N/A ± N/A | N/A ± N/A |

Data are given as mean ± SEM

We find that already early in development some SPNs form local inhibitory synapses with each other with a low incidence initially but this progressively increases with age (P3-6: 2.3%, P9-12: 6.9% and P21+: 12.2%, n=9/379, 28/408 and 24/197, P3-6 vs P21+ p=0.0431, Fisher exact test, **Fig. 5H**). Several observations can be made. Firstly, the earliest synaptic connections at P3-6 can be found from D1 SPNs to D1 and D2 SPNs (D1 to D1: 3.0% and D1 to D2: 3.3%; n=2/67 and n=2/60, **Fig. 5I**) potentially a reflection of their earlier birthdate (Marchand & Lajoie, 1986; van der Kooy & Fishell, 1987; Kelly *et al.*, 2018) as we did not detect any synaptic connections from D2 SPN. Secondly, of all synaptic connections that were found at younger ages, a subset was also connected through gap junctions (P3-6: 33.3% and P9-12: 14.3%). Thirdly, the majority of synaptic connections were unidirectional (P3-6: 88.89%, P9-12: 75.0% and P21+: 68.42%). Fourthly, already at P9-12 the relative incidence and biases in synaptic connectivity seen in adulthood between D1 and D2 SPNs (Planert *et al.*, 2010) can already be found and include a high incidence of synaptic connections between D2 SPNs (P9-P12; D1 to D1: 6.5%, D1 to D2: 5.1%, D2 to D2: 12.8% and D2 to D1: 5.3% n=3/46, n=4/78, 13/102 and 4/76, **Fig. 5I**). Indeed, these relative biases in synaptic connections are maintained or further strengthened by P21+ (D1 to D1: 7.1%, D1 to D2: 8.0%, D2 to D2: 21.1% and D2 to D1: 13.9% n=1/14, n=2/25, 12/57 and 5/36, **Fig. 5I**) and are consistent with those previously described (Taverna *et al.*, 2008; Planert *et al.*, 2010). Not only are the D2-D2 SPN connections most numerous at P9-12 and P21+ (P9-12: 12.8% and P21+: 21.1%) they also form the strongest mutual synaptic connections (P9-12: 0.97 ± 0.32 mV and P21+: 0.72 ± 0.28 mV, **Table 4**) as found by others also (Planert *et al.*, 2010). We do not find significant differences in other properties of the synaptic connections between the different SPN types (Taverna *et al.*, 2008) or across the age ranges (**Table 4**). Lastly, we looked at the short-term plastic properties of the inhibitory synaptic connections and find that these become depressing as the synapses mature suggesting a progressive increase in release probability (F(1, 132)=14.85, p=0.00018, **Fig. 5J**).

**Table 4:**
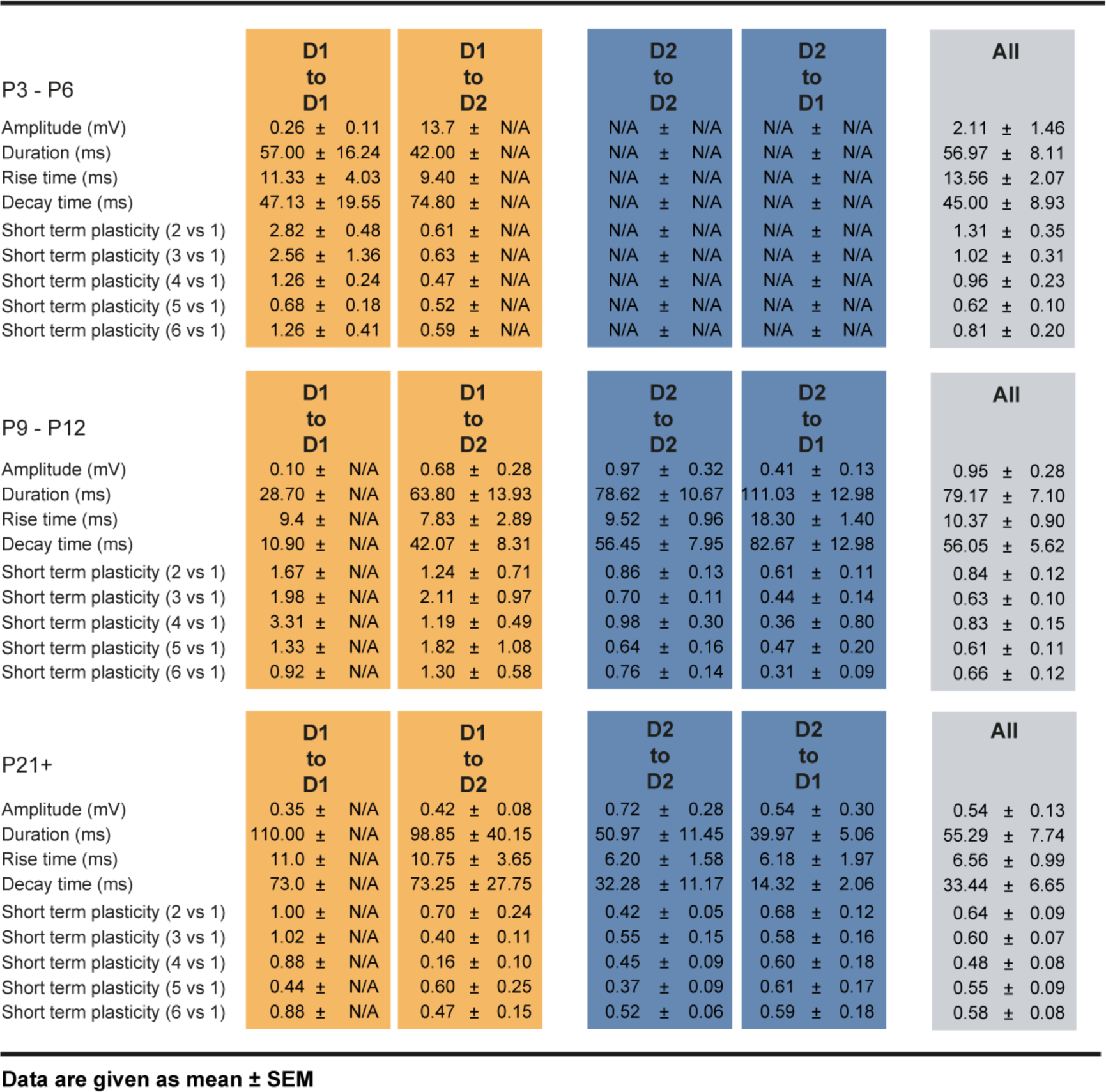
Properties of unitary GABAergic synapses between D1 and D2 SPNs

In conclusion, these results show that as the striatal circuit matures, both D1 and D2 SPNs gradual replace their gap junctions with precise local inhibitory synaptic connections. Indeed, they reveal that biases in inhibitory synaptic connectivity, for example the high incidence of local inhibitory synaptic connections between D2 SPNS, are already apparent in the second postnatal week.

**Figure 5:**
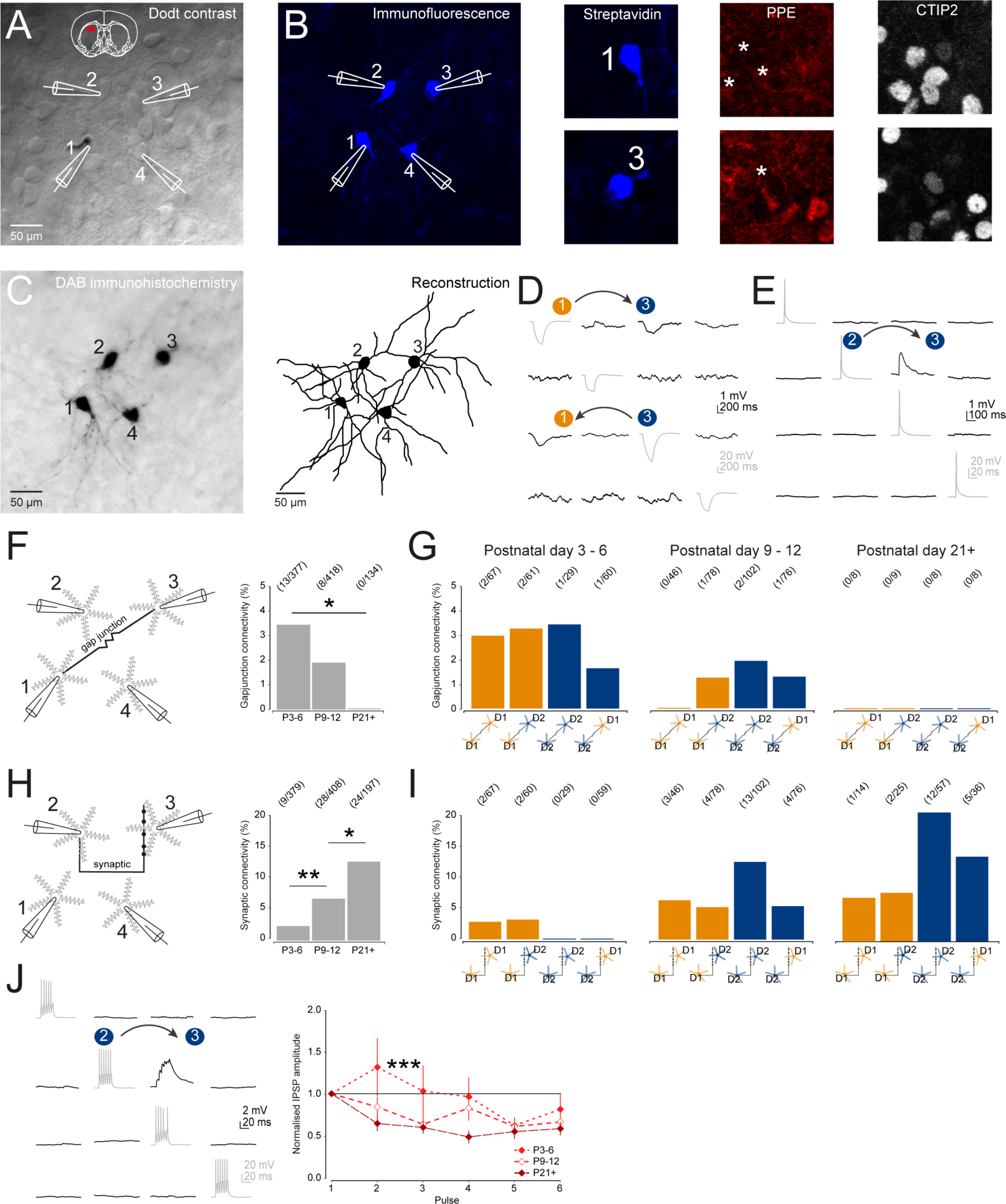
Development of local gap junctions and inhibitory synaptic connections between SPNs. (**A**) Dodt contrast image of recording configuration consisting of four simultaneously recorded SPNs. (**B**) Posthoc immunocytochemistry of recorded neurons using antibodies against streptavidin, PPE and CTIP2 allowed for classification of neurons as D1 or D2 SPNs. (**C**) Subsequently the slices were processed for DAB immunohistochemistry to label SPNs (left) and reveal fine detailed dendritic structures allowing reconstruction of SPNs (right). (**D**) Hyperpolarizing current steps revealed the presence of potential gap junctions connecting recorded SPNs. Note the presence of bidirectional gap junctions between SPN no.1 and no.3. (**E**) Suprathreshold current injections elicited action potentials in recorded SPNs and revealed potential synaptic connections in other simultaneously recorded SPNs. Note the presence of a unidirectional synaptic connection from SPN no.2 to SPN no.3. (**F**) Diagram of experimental setup to test for potential gap junctions between SPNs (left). Barplot showing a significant decrease in the incidence of detected gap junctions as the SPNs mature (right). (**G**) Barplots of gap junction incidence between the SPN connection groups across the age ranges. Note the relative uniform incidence of gap junctions in all SPN groups at P3-6 followed by a progressive reduction and absence of detected gap junctions at P21+. (**H**) Diagram of experimental setup to test for synaptic connections between SPNs (left). Barplot showing a progressive and significant increase in the incidence of detected synaptic connections as the SPNs mature (right). (**I**) Synaptic connectivity between the SPN connection across the age ranges. Note the earliest appearance of synaptic connections from D1 SPNs at P3-6. At P9-12 synaptic connections between all types of SPNs have appeared and relative biases seen at P21+ are already apparent at this age. (**J**) Trains of action potentials to presynaptic neurons (6 pulses at 20 Hz) revealed that as SPNs mature the short-term plasticity of their synapses changes from facilitating to depressing.

## Discussion

In this paper we describe the developmental trajectory of identified D1 and D2 SPNs from the earliest postnatal period (P3-6) into adulthood (P21+). We find that young D1 SPNs are electrically more mature in the first postnatal week and differences in intrinsic excitability of D1 and D2 SPN become apparent in the second postnatal week and are maintained into adulthood. Secondly, both SPNs exhibit a radial dendritic morphology from birth that develops in complexity in parallel across the age ranges. Thirdly, both excitatory and inhibitory synaptic synapses are functional in the first postnatal week with the inhibitory synaptic inputs exhibiting a comparatively prolonged maturation. Indeed, most SPNs receive long-range excitatory synaptic inputs from cortex during the first postnatal week, which progressively strengthen through postsynaptic glutamate receptor changes. Initially, SPNs communicate locally through gap junctions but increasingly via inhibitory synaptic connections in the second postnatal week. Interestingly, the relative biases in inhibitory connections between D1 and D2 SPNs are already apparent in the second postnatal week and are maintained into adulthood. Overall, we find that the developmental trajectory is similar for both the D1 and D2 SPNs and that many of the mature neuronal and circuit properties are apparent by the second postnatal week suggesting a major role for intrinsic programs in guiding their development.

### Intrinsic cellular properties of D1 and D2 SPNs

We find a progressive development of both the intrinsic electrophysiological and morphological properties of the D1 and D2 SPNs. Both are able to generate small immature action potentials during the first few postnatal days and both undergo a progressive decrease in their input resistance, a hyperpolarization of their resting membrane potential (Lieberman *et al.*, 2018) and an increase in their firing frequency (Peixoto *et al.*, 2016). However, some differences were observed. Initially, the size and duration of the action potentials were more mature for the D1 SPNs, possibly as a result from their suggested earlier birthdate (Marchand & Lajoie, 1986; van der Kooy & Fishell, 1987; Kelly *et al.*, 2018). Secondly, many of the differences in electrical properties of SPNs, such as the comparatively more depolarized resting membrane potential, higher input resistance and higher firing frequency of D2 SPNs, are already apparent in the second postnatal week and are maintained into adulthood (Gertler *et al.*, 2008; Peixoto *et al.*, 2016; Lieberman *et al.*, 2018). Morphologically we find that across the age ranges there is a substantial elaboration of the SPN dendritic arbor concomitant with an increase in dendritic spine density occurring in parallel for the D1 and D2 SPNs. We could not morphologically distinguish the D1 and D2 SPNs at any of the ages suggesting that previously described differences (Gertler *et al.*, 2008; Kozorovitskiy *et al.*, 2012) might be due to specifics of age, mouse line or methodology (Bagetta *et al.*, 2011; Kramer *et al.*, 2011; Chan *et al.*, 2012; Nelson *et al.*, 2012). We do find a progressive increase in dendritic spine density consistent with previous studies (Sharpe & Tepper, 1998; Tepper *et al.*, 1998), but an overall lower spine density as compared to 2-photon imaging and serial section EM studies (Ingham *et al.*, 1998; Day *et al.*, 2006; Kozorovitskiy *et al.*, 2012) likely due to our use of DAB immunohistochemistry and occlusion of spines on the top and bottom of dendrites by the DAB reaction product (Ingham *et al.*, 1998; Day *et al.*, 2006).

### Functional excitatory synaptic inputs to D1 and D2 SPNs

We find that already soon after birth excitatory inputs onto SPNs are functional and are able to depolarize both D1 and D2 SPNs. Whereas the frequency of mEPSCs at P3-6 is close to that observed in adulthood we see a dramatic increase in mEPSC amplitude for both D1 and D2 SPNs from the first to second postnatal week, consistent with previous observations (Dehorter *et al.*, 2009; Peixoto *et al.*, 2016) and suggestive of postsynaptic changes. Indeed, in our recordings of electrically evoked corticostriatal responses we find that the majority of D1 and D2 SPNs have functional synapses at P3-6 and both exhibit similar increases in amplitude (Day *et al.*, 2006). This increase in amplitude correlates with a relative increase in AMPA receptor-mediated currents and concomitant decrease in the NMDA/AMPA receptor ratio seen in both D1 and D2 SPNs (Colwell *et al.*, 1998; Hurst *et al.*, 2001; Peixoto *et al.*, 2016). Indeed, such changes in glutamate receptors in this developmental period have been shown to occur at corticostriatal synapses (Dehorter *et al.*, 2009; Peixoto *et al.*, 2016) and in many other brain regions (Lohmann & Kessels, 2014). Interestingly, these sets of experiments reveal a stable electrically evoked corticostriatal EPSP amplitude in both D1 and D2 SPNs across developmental time (Tepper *et al.*, 1998), suggesting that dynamic changes in SPNs, such as decreasing input resistance and increased excitatory postsynaptic currents, are balanced and result in a stable excitatory drive to SPNs. We were surprised to see a relatively stable mEPSC frequency despite reported increases in the density of asymmetric glutamatergic synapses during this period (Butler *et al.*, 1998; Sharpe & Tepper, 1998; Tepper *et al.*, 1998) and previous reported increases in EPSC frequency (Peixoto *et al.*, 2016). Possible explanations for our observations and others (Dehorter *et al.*, 2011) might be biases towards recording mEPSCs mediated by axo-dendritic synapses instead of axo-spinous synapses, as the density of axo-dendritic synapses remain stable during development (Sharpe & Tepper, 1998), or could result from a dynamic interplay between synapse number and release probability where increases in synapse number are balanced by decreases in release probability. Indeed, the observed changes in the short-term plasticity at D1 SPN corticostriatal synapses are consistent with a decrease in their release probability (Choi & Lovinger, 1997). One of the other major contributor to our mEPSC measurements originates from excitatory afferents coming from the heterogeneous population of thalamic nuclei (Ellender *et al.*, 2013; Smith *et al.*, 2014) which project to the striatum and are thought to arrive relatively early in development (Nakamura *et al.*, 2005). When and how these afferents develop and innervate striatal SPNs remains to be investigated.

### Local connections between D1 and D2 SPNs

The activity of striatal SPNs and their response to excitatory input is modulated by inhibition coming from local collaterals from neighboring SPNs (Somogyi *et al.*, 1981; Bolam & Izzo, 1988; Taverna *et al.*, 2008; Planert *et al.*, 2010; Cepeda *et al.*, 2013) and striatal interneurons (Tepper & Plenz, 2006; Ponzi & Wickens, 2010). Measurements of mIPSCs reveal that both D1 and D2 SPNs already receive some inhibitory input in the first postnatal week. The initial frequency of these events is lower than that seen for mEPSCs but progressively increase in the second and third postnatal week as previously observed (Dehorter *et al.*, 2011). This would suggest that GABAergic synapse density increases well into the third postnatal week. We also observed a dramatic increase in mIPSC amplitude from the first to second postnatal week consistent with a rapid maturation as a result of an increased number (Nusser *et al.*, 1997) and/or changed subunit composition of postsynaptic GABA receptors (Farrant & Nusser, 2005; Arama *et al.*, 2015). To investigate when and how SPNs start communicating with each other in the striatum and whether the number of synaptic connections does increase we performed simultaneous quadruple whole-cell patch-clamp recordings of SPNs. We find that both D1 and D2 SPNs are mainly connected through gap junctions in the first postnatal week, but both the incidence of gap junctions and their coupling coefficient rapidly decrease and we do not observe gap junctions in adulthood, consistent with previous electrophysiological (Venance *et al.*, 2004; Yu *et al.*, 2012) and dye coupling experiments (Tepper *et al.*, 1998). These initial gap junctions could facilitate synchronization of SPN activity (Venance *et al.*, 2004; Hestrin & Galarreta, 2005) and the establishment of synaptic connections (Yu *et al.*, 2012). The first inhibitory synaptic connections can be detected in the first postnatal week and come from D1 SPNs but in the second postnatal week both D1 and D2 SPNs form inhibitory synaptic connections with each other. Interestingly, the relative biases in connectivity (e.g. the high connectivity probability between D2 SPNs) seen in adulthood in this study and by others (Planert *et al.*, 2010) are already established in the second postnatal week prior to when sensory inputs from afferent structures is thought to have arrived (Tobach *et al.*, 1971; Ko *et al.*, 2013; Mowery *et al.*, 2015; 2016; Mowery *et al.*, 2017).

In conclusion we find that the intrinsic cellular and circuit properties of the D1-expressing direct pathway and the D2-expressing indirect pathway SPNs exhibit many developmental similarities. Nonetheless, mature D1 and D2 SPNs exhibit distinct electrical properties and form precise local inhibitory connections, which we show are already apparent in the second postnatal week before sensory input from afferent structures is thought to have arrived (Tobach *et al.*, 1971; Ko *et al.*, 2013; Mowery *et al.*, 2015; 2016; Mowery *et al.*, 2017). This leaves scope for factors such as intrinsic genetic programs (Arlotta *et al.*, 2008), spontaneous early network patterns (Khazipov *et al.*, 2004; Hanganu-Opatz, 2010; Dehorter *et al.*, 2011; Ackman *et al.*, 2012) and neuromodulators (Kozorovitskiy *et al.*, 2015; Lieberman *et al.*, 2018) in shaping the development of the D1 and D2 SPNs and might inform future investigations of striatal formation in neurodevelopmental disorders (Graybiel & Rauch, 2000; Del Campo *et al.*, 2011; Langen *et al.*, 2011; McNaught & Mink, 2011; Shepherd, 2013; Albin, 2018).

## Methods and Materials

### Animals

All experiments were carried out on C57/BL6 wildtype and heterozygous D1-GFP or D2-GFP mice. The D1-GFP or D2-GFP BAC transgenic mice report subtypes of the dopamine receptor, either D1 or D2, by the presence of GFP (Mutant Mouse Regional Resource Centers, MMRRC). Details of the mice and the methods of BAC mice production have been published (Gong *et al.*, 2003) and can be found on the GENSAT website [GENSAT (2009) The Gene Expression Nervous System Atlas (GENSAT) Project. In: NINDS, Contracts N01NS02331 and HHSN271200723701C, The Rockefeller University (New York), http://www.gensat.org/index.html. The BAC transgenic mice were backcrossed to a C57/BL6 background over 10+ generations prior to use and kept as a heterozygous mouse line to avoid published issues using these transgenic lines (Bagetta *et al.*, 2011; Kramer *et al.*, 2011; Chan *et al.*, 2012; Nelson *et al.*, 2012). All mice were bred, housed and used in accordance with the UK Animals (Scientific Procedures) Act (1986).

### Slice preparation and recording conditions

Acute striatal slices were made from postnatal animals between postnatal 3-6, postnatal day 9-12 or postnatal day 21 and later. Animals were anaesthetized with isoflurane and then decapitated. Coronal 350-400 μm slices were cut using a vibrating microtome (Microm HM650V). Slices were prepared in artificial cerebrospinal fluid (aCSF) containing (in mM): 65 Sucrose, 85 NaCl, 2.5 KCl, 1.25 NaH_2_PO_4_, 7 MgCl_2_, 0.5 CaCl_2_, 25 NaHCO_3_ and 10 glucose, pH 7.2-7.4, bubbled with carbogen gas (95% O_2_ / 5% CO_2_). Slices were immediately transferred to a storage chamber containing aCSF (in mM): 130 NaCl, 3.5 KCl, 1.2 NaH_2_PO_4_, 2 MgCl_2_, 2 CaCl_2_, 24 NaHCO_3_ and 10 glucose, pH 7.2-7.4, at 32 ^°^C and bubbled with carbogen gas until used for recording. Striatal slices were transferred to a recording chamber and continuously superfused with aCSF bubbled with carbogen gas with the same composition as the storage solution (32 ^°^C and perfusion speed of 2 ml/min). Whole-cell current-clamp recordings were performed using glass pipettes, pulled from standard wall borosilicate glass capillaries and containing for whole-cell current clamp (in mM): 110 potassium gluconate, 40 HEPES, 2 ATP-Mg, 0.3 Na-GTP, 4 NaCl and 4 mg/ml biocytin (pH 7.2-7.3; osmolarity, 290-300 mosmol/l) and for whole-cell voltage-clamp (in mM): 120 cesium gluconate, 40 HEPES, 4 NaCl, 2 ATP-Mg, 0.3 Na-GTP, 0.2 QX-314 and 4 mg/ml biocytin (pH 7.2–7.3; osmolarity, 290-300 mosmol/L). Recording of gap junction and synaptic connections between SPNs used an intracellular solution with high internal [Cl^-^] containing (in mM): 105 potassium gluconate, 30 KCl, 10 HEPES, 4 ATP-Mg, 0.3 Na-GTP, and 4 mg/ml biocytin (pH 7.2-7.3; osmolarity, 290-300 mosmol/l). Recordings were made using Multiclamp 700B amplifiers and filtered at 4kHz and acquired using an InstruTECH ITC-18 analog/digital board and WinWCP software (University of Strathclyde, RRID:SCR_014713) at 10 kHz.

### Stimulation and recording protocols

Hyperpolarizing and depolarizing current steps were used to assess the intrinsic properties of the recorded SPNs including input resistance and spike threshold (using 1-10 pA incremental current steps) as well as the properties of action potentials (amplitude, frequency and duration). Currents steps were for P3-6: −50pA to +50pA, for P9-12: −100pA to +100pA and for P21+: −500pA to +500 pA. These ranges of currents were chosen to allow sufficient depolarization of SPNs taking in consideration changes in input resistance and observations of depolarization block in SPNs. Miniature EPSCs and IPSCs were recorded in 5 min traces from SPNs held at respectively −75 mV and 0 mV in the presence of 1 μM TTX in the recording aCSF. Activation of excitatory cortical afferents was performed via a bipolar stimulating electrode (FHC Inc., USA) placed in the external capsule in the presence of blockers of inhibitory GABAergic transmission including the GABA_A_-receptor antagonist SR95531 (1 μM) and the GABA_B_-receptor antagonist CGP52432 (2 μM). Afferents were activated every 5s with up to 20 repetitions and excitatory postsynaptic currents (EPSCs) and excitatory postsynaptic potentials (EPSPs) were recorded from the patched SPNs. Evoked EPSCs were recorded in whole-cell voltage-clamp mode at a holding potential near −75 mV and evoked EPSPs in whole-cell current-clamp mode at resting membrane potential. Trains of stimulation consisted of nine pulses given at 20 Hz and trains were repeated every 30 s up to 5 times. Combined AMPA and NMDA receptor-mediated currents were recorded from SPNs held at +50 mV. AMPA receptor-mediated currents were recorded after a 5-10 min wash-in of d-AP5 (50 μM). Local gap junctions between SPNs were examined by delivering hyperpolarizing current injections (200 ms, P3-6: −20 pA, P9-12: −100 pA and P21+: −200 pA) to each patched SPN sequentially, whilst simultaneously recording the membrane voltage of the other SPNs. Local SPN inhibitory synaptic connectivity was examined by delivering brief (∼60 ms) suprathreshold current injections (P3-6: +50 pA, P9-12: +150 pA and P21+: +400 pA) or brief trains of current injections (6 pulses, 30 ms, P3-6: +80 pA, P9-12: +200 pA and P21+: +500 pA at 20 Hz) to each patched SPN sequentially, whilst simultaneously recording the membrane voltage of the other SPNs. These protocols were repeated 20-30 times.

### Analysis of recordings

Data were analyzed offline using custom written programs in Igor Pro (Wavemetrics, RRID:SCR_000325). The input resistance was calculated from the observed membrane potential change after hyperpolarizing the membrane potential with a set current injection. The spike threshold was the membrane voltage at which the SPN generated an action potential. mEPSCs and mIPSCs were detected as downward and upward deflections of more than 2 standard deviations (SD) above baseline and more than 10 ms duration in 5 min traces which were lowpass filtered at 50Hz. Evoked EPSCs, EPSPs and IPSPs were defined as upward or downward deflections of more than 2 standard deviations (SD) on average synaptic responses generated after filtering original traces (0.1 Hz high pass filter and 500 Hz low pass filter) and averaging over all traces and used for analysis of synaptic properties (peak amplitude, duration, rise time (between 20% and 80% from peak) and decay time). The short-term plastic properties of excitatory synapses and unitary inhibitory synapses between SPNs were analyzed by taking the amplitude of each EPSP/IPSP during train stimulation and dividing this by the amplitude of the first response. NMDA/AMPA receptor ratio was calculated from an average trace of combined AMPA and NMDA receptor-mediated current as well as the pharmacologically isolated AMPA receptor-mediated current. The average AMPA receptor-mediated current trace was subtracted from the combined AMPA and NMDA receptor-mediated current trace to obtain the NMDA receptor-mediated current. Peak amplitude NMDA receptor-mediated current was divided by peak amplitude AMPA receptor-mediated current to obtain the NMDA/AMPA ratio. The presence of gap junctions was assessed by averaging the 20-30 sweeps of hyperpolarizing current injections and observing a significant downward deflection of more than 2 standard deviations (SD) from baseline. The coupling coefficient (CC) was obtained by dividing the amplitude of the low-frequency voltage change in the receiver SPN to that in the driver SPN. The junctional conductance (*G*_j_) was estimated from *R*_input_ and CC (Venance *et al.*, 2004): *G*_j1-2_=R_input1_ x CC_1-2_/((R_input1_ x R_input2_)-(R_input1_ x CC_1-2_)^2^), where Rinput1 and Rinput2 are the R_input_ values of the injected and receiving SPNs, respectively and CC_1-2_ the CC between the injected and receiving SPNS.

### Histological analyses

Following whole-cell patch-clamp recordings the brain slices were fixed in 4% paraformaldehyde in 0.1 M phosphate buffer (PB; pH 7.4). Biocytin-filled neurons were visualized by incubating sections in 1:10,000 streptavidin AlexaFluor405-conjugated antibodies (ThermoFisher Scientific, Cat#S32351). Visualized neurons were labeled for chicken ovalbumin upstream promoter transcription-factor interacting protein-2 (CTIP2, 1:1000, rat, Abcam, Cat#ab14865, RRID:AB_2064130) and pre-proenkephalin (PPE, 1:1000, rabbit, LifeSpan Biosciences, Cat#LS-C23084, RRID:AB_902714) in PBS containing 0.3% Triton X-100 (PBS-Tx) overnight at 4^°^C followed by incubation with goat-anti-rat AlexaFluor647 (1:500; ThermoFisher Scientific, CAT#:A-21247, RRID:AB_141778) and goat-anti-rabbit AlexaFluor555 (1:500; ThermoFisher Scientific, CAT#:A-21429, RRID:AB_2535850) secondary antibodies in 0.3% PBS-Tx for 2 h at RT for SPN classification. PPE antibody staining was facilitated through antigen retrieval by heating sections at 80^°^ C in 10 mM sodium citrate (pH 6.0) for approximately 30-60 min prior to incubation with adding PPE primary antibody. After classification of SPN type slices were washed 3 times in PBS and processed for DAB immunohistochemistry using standard procedures. Fluorescence images were captured with a LSM 710 confocal microscope using ZEN software (Zeiss, RRID:SCR_013672). DAB-immunoreactive neurons were visualized on a brightfield microscope and were reconstructed and analyzed using Neurolucida and Neuroexplorer software (MBF Bioscience, RRID:SCR_001775).

### Statistics

All data are presented as means ± SEM. Statistical tests were all two-tailed and performed using SPSS 17.0 (IBM SPSS statistics, RRID:SCR_002865) or GraphPad Prism version 5.0 (GraphPad software, RRID:SCR_002798). Synaptic connectivity ratios were compared with Fisher’s Exact test. Continuous data were assessed for normality and appropriate parametric (ANOVA, paired t-test and unpaired t-test) or non-parametric Mann-Whitney U statistical tests were applied (* p<0.05, ** p<0.01, *** p<0.001).

## Acknowledgements

We would like to thank Peter Somogyi, Peter Magill and Colin Akerman for generously providing access to equipment and Ben Micklem for providing technical assistance. TJE was supported by an MRC Career Development Award (MR/M009599/1) and AMD by an Imperial College research bursary.

## References

Ackman, J.B., Burbridge, T.J. & Crair, M.C. (2012) Retinal waves coordinate patterned activity throughout the developing visual system. Nature, 490, 219–225.

Albin, R.L. (2018) Tourette syndrome: a disorder of the social decision-making network. Brain, 141, 332–347.

Arama, J., Abitbol, K., Goffin, D., Fuchs, C., Sihra, T.S., Thomson, A.M. & Jovanovic, J.N. (2015) GABAA receptor activity shapes the formation of inhibitory synapses between developing medium spiny neurons. Frontiers in cellular neuroscience, 9, 290.

Arlotta, P., Molyneaux, B.J., Jabaudon, D., Yoshida, Y. & Macklis, J.D. (2008) Ctip2 controls the differentiation of medium spiny neurons and the establishment of the cellular architecture of the striatum. J Neurosci, 28, 622–632.

Bagetta, V., Picconi, B., Marinucci, S., Sgobio, C., Pendolino, V., Ghiglieri, V., Fusco, F.R., Giampa, C. & Calabresi, P. (2011) Dopamine-dependent long-term depression is expressed in striatal spiny neurons of both direct and indirect pathways: implications for Parkinson’s disease. J Neurosci, 31, 12513–12522.

Belousov, A.B. & Fontes, J.D. (2013) Neuronal gap junctions: making and breaking connections during development and injury. Trends in neurosciences, 36, 227–236.

Bolam, J.P. & Izzo, P.N. (1988) The postsynaptic targets of substance P-immunoreactive terminals in the rat neostriatum with particular reference to identified spiny striatonigral neurons. Experimental brain research. Experimentelle Hirnforschung, 70, 361–377.

Buchwald, N.A., Price, D.D., Vernon, L. & Hull, C.D. (1973) Caudate intracellular response to thalamic and cortical inputs. Experimental neurology, 38, 311–323.

Butler, A.K., Uryu, K. & Chesselet, M.F. (1998) A role for N-methyl-D-aspartate receptors in the regulation of synaptogenesis and expression of the polysialylated form of the neural cell adhesion molecule in the developing striatum. Developmental neuroscience, 20, 253–262.

Cepeda, C., Galvan, L., Holley, S.M., Rao, S.P., Andre, V.M., Botelho, E.P., Chen, J.Y., Watson, J.B., Deisseroth, K. & Levine, M.S. (2013) Multiple sources of striatal inhibition are differentially affected in Huntington’s disease mouse models. J Neurosci, 33, 7393–7406.

Chan, C.S., Peterson, J.D., Gertler, T.S., Glajch, K.E., Quintana, R.E., Cui, Q., Sebel, L.E., Plotkin, J.L., Shen, W., Heiman, M., Heintz, N., Greengard, P. & Surmeier, D.J. (2012) Strain-specific regulation of striatal phenotype in Drd2-eGFP BAC transgenic mice. J Neurosci, 32, 9124–9132.

Choi, S. & Lovinger, D.M. (1997) Decreased probability of neurotransmitter release underlies striatal long-term depression and postnatal development of corticostriatal synapses. Proceedings of the National Academy of Sciences of the United States of America, 94, 2665–2670.

Colwell, C.S., Cepeda, C., Crawford, C. & Levine, M.S. (1998) Postnatal development of glutamate receptor-mediated responses in the neostriatum. Developmental neuroscience, 20, 154–163.

Day, M., Wang, Z., Ding, J., An, X., Ingham, C.A., Shering, A.F., Wokosin, D., Ilijic, E., Sun, Z., Sampson, A.R., Mugnaini, E., Deutch, A.Y., Sesack, S.R., Arbuthnott, G.W. & Surmeier, D.J. (2006) Selective elimination of glutamatergic synapses on striatopallidal neurons in Parkinson disease models. Nature neuroscience, 9, 251–259.

Day, M., Wokosin, D., Plotkin, J.L., Tian, X. & Surmeier, D.J. (2008) Differential excitability and modulation of striatal medium spiny neuron dendrites. J Neurosci, 28, 11603–11614.

Dehorter, N., Guigoni, C., Lopez, C., Hirsch, J., Eusebio, A., Ben-Ari, Y. & Hammond, C. (2009) Dopamine-deprived striatal GABAergic interneurons burst and generate repetitive gigantic IPSCs in medium spiny neurons. J Neurosci, 29, 7776–7787.

Dehorter, N., Michel, F.J., Marissal, T., Rotrou, Y., Matrot, B., Lopez, C., Humphries, M.D. & Hammond, C. (2011) Onset of Pup Locomotion Coincides with Loss of NR2C/D-Mediated Cortico-Striatal EPSCs and Dampening of Striatal Network Immature Activity. Frontiers in cellular neuroscience, 5, 24.

Del Campo, N., Chamberlain, S.R., Sahakian, B.J. & Robbins, T.W. (2011) The roles of dopamine and noradrenaline in the pathophysiology and treatment of attention-deficit/hyperactivity disorder. Biological psychiatry, 69, e145–157.

Ellender, T.J., Harwood, J., Kosillo, P., Capogna, M. & Bolam, J.P. (2013) Heterogeneous properties of central lateral and parafascicular thalamic synapses in the striatum. The Journal of physiology, 591, 257–272.

Farrant, M. & Nusser, Z. (2005) Variations on an inhibitory theme: phasic and tonic activation of GABA(A) receptors. Nature reviews, 6, 215–229.

Gerfen, C.R., Engber, T.M., Mahan, L.C., Susel, Z., Chase, T.N., Monsma, F.J., Jr. & Sibley, D.R. (1990) D1 and D2 dopamine receptor-regulated gene expression of striatonigral and striatopallidal neurons. Science, 250, 1429–1432.

Gertler, T.S., Chan, C.S. & Surmeier, D.J. (2008) Dichotomous anatomical properties of adult striatal medium spiny neurons. J Neurosci, 28, 10814–10824.

Gong, S., Zheng, C., Doughty, M.L., Losos, K., Didkovsky, N., Schambra, U.B., Nowak, N.J., Joyner, A., Leblanc, G., Hatten, M.E. & Heintz, N. (2003) A gene expression atlas of the central nervous system based on bacterial artificial chromosomes. Nature, 425, 917–925.

Graybiel, A.M., Aosaki, T., Flaherty, A.W. & Kimura, M. (1994) The basal ganglia and adaptive motor control. Science, 265, 1826–1831.

Graybiel, A.M. & Rauch, S.L. (2000) Toward a neurobiology of obsessive-compulsive disorder. Neuron, 28, 343–347.

Grillner, S., Hellgren, J., Menard, A., Saitoh, K. & Wikstrom, M.A. (2005) Mechanisms for selection of basic motor programs--roles for the striatum and pallidum. Trends in neurosciences, 28, 364–370.

Hanganu-Opatz, I.L. (2010) Between molecules and experience: role of early patterns of coordinated activity for the development of cortical maps and sensory abilities. Brain Res Rev, 64, 160–176.

Hestrin, S. & Galarreta, M. (2005) Electrical synapses define networks of neocortical GABAergic neurons. Trends in neurosciences, 28, 304–309.

Hurst, R.S., Cepeda, C., Shumate, L.W. & Levine, M.S. (2001) Delayed postnatal development of NMDA receptor function in medium-sized neurons of the rat striatum. Developmental neuroscience, 23, 122–134.

Ingham, C.A., Hood, S.H., Taggart, P. & Arbuthnott, G.W. (1998) Plasticity of synapses in the rat neostriatum after unilateral lesion of the nigrostriatal dopaminergic pathway. J Neurosci, 18, 4732–4743.

Kelly, S.M., Raudales, R., He, M., Lee, J.H., Kim, Y., Gibb, L.G., Wu, P., Matho, K., Osten, P., Graybiel, A.M. & Huang, Z.J. (2018) Radial Glial Lineage Progression and Differential Intermediate Progenitor Amplification Underlie Striatal Compartments and Circuit Organization. Neuron.

Kemp, J.M. & Powell, T.P. (1971) The site of termination of afferent fibres in the caudate nucleus. Philos Trans R Soc Lond B Biol Sci, 262, 413–427.

Khazipov, R., Sirota, A., Leinekugel, X., Holmes, G.L., Ben-Ari, Y. & Buzsaki, G. (2004) Early motor activity drives spindle bursts in the developing somatosensory cortex. Nature, 432, 758–761.

Ko, H., Cossell, L., Baragli, C., Antolik, J., Clopath, C., Hofer, S.B. & Mrsic-Flogel, T.D. (2013) The emergence of functional microcircuits in visual cortex. Nature, 496, 96–100.

Kozorovitskiy, Y., Peixoto, R., Wang, W., Saunders, A. & Sabatini, B.L. (2015) Neuromodulation of excitatory synaptogenesis in striatal development. Elife, 4.

Kozorovitskiy, Y., Saunders, A., Johnson, C.A., Lowell, B.B. & Sabatini, B.L. (2012) Recurrent network activity drives striatal synaptogenesis. Nature, 485, 646–650.

Kramer, P.F., Christensen, C.H., Hazelwood, L.A., Dobi, A., Bock, R., Sibley, D.R., Mateo, Y. & Alvarez, V.A. (2011) Dopamine D2 receptor overexpression alters behavior and physiology in Drd2-EGFP mice. J Neurosci, 31, 126–132.

Kravitz, A.V., Freeze, B.S., Parker, P.R., Kay, K., Thwin, M.T., Deisseroth, K. & Kreitzer, A.C. (2010) Regulation of parkinsonian motor behaviours by optogenetic control of basal ganglia circuitry. Nature, 466, 622–626.

Langen, M., Kas, M.J., Staal, W.G., van Engeland, H. & Durston, S. (2011) The neurobiology of repetitive behavior: of mice. Neurosci Biobehav Rev, 35, 345–355.

Lieberman, O.J., McGuirt, A.F., Mosharov, E.V., Pigulevskiy, I., Hobson, B.D., Choi, S., Frier, M.D., Santini, E., Borgkvist, A. & Sulzer, D. (2018) Dopamine Triggers the Maturation of Striatal Spiny Projection Neuron Excitability during a Critical Period. Neuron, 99, 540–554 e544.

Lohmann, C. & Kessels, H.W. (2014) The developmental stages of synaptic plasticity. The Journal of physiology, 592, 13–31.

Marchand, R. & Lajoie, L. (1986) Histogenesis of the striopallidal system in the rat. Neurogenesis of its neurons. Neuroscience, 17, 573–590.

McNaught, K.S. & Mink, J.W. (2011) Advances in understanding and treatment of Tourette syndrome. Nat Rev Neurol, 7, 667–676.

Monyer, H., Burnashev, N., Laurie, D.J., Sakmann, B. & Seeburg, P.H. (1994) Developmental and regional expression in the rat brain and functional properties of four NMDA receptors. Neuron, 12, 529–540.

Mowery, T.M., Kotak, V.C. & Sanes, D.H. (2015) Transient Hearing Loss Within a Critical Period Causes Persistent Changes to Cellular Properties in Adult Auditory Cortex. Cereb Cortex, 25, 2083–2094.

Mowery, T.M., Kotak, V.C. & Sanes, D.H. (2016) The onset of visual experience gates auditory cortex critical periods. Nature communications, 7, 10416.

Mowery, T.M., Penikis, K.B., Young, S.K., Ferrer, C.E., Kotak, V.C. & Sanes, D.H. (2017) The Sensory Striatum Is Permanently Impaired by Transient Developmental Deprivation. Cell reports, 19, 2462–2468.

Nakamura, K., Hioki, H., Fujiyama, F. & Kaneko, T. (2005) Postnatal changes of vesicular glutamate transporter (VGluT)1 and VGluT2 immunoreactivities and their colocalization in the mouse forebrain. The Journal of comparative neurology, 492, 263–288.

Nelson, A.B., Hang, G.B., Grueter, B.A., Pascoli, V., Luscher, C., Malenka, R.C. & Kreitzer, A.C. (2012) A comparison of striatal-dependent behaviors in wild-type and hemizygous Drd1a and Drd2 BAC transgenic mice. J Neurosci, 32, 9119–9123.

Nusser, Z., Cull-Candy, S. & Farrant, M. (1997) Differences in synaptic GABA(A) receptor number underlie variation in GABA mini amplitude. Neuron, 19, 697–709.

Peixoto, R.T., Wang, W., Croney, D.M., Kozorovitskiy, Y. & Sabatini, B.L. (2016) Early hyperactivity and precocious maturation of corticostriatal circuits in Shank3B(-/-) mice. Nature neuroscience, 19, 716–724.

Planert, H., Szydlowski, S.N., Hjorth, J.J., Grillner, S. & Silberberg, G. (2010) Dynamics of synaptic transmission between fast-spiking interneurons and striatal projection neurons of the direct and indirect pathways. J Neurosci, 30, 3499–3507.

Ponzi, A. & Wickens, J. (2010) Sequentially switching cell assemblies in random inhibitory networks of spiking neurons in the striatum. J Neurosci, 30, 5894–5911.

Seeburg, P.H. (1993) The TINS/TiPS Lecture. The molecular biology of mammalian glutamate receptor channels. Trends in neurosciences, 16, 359–365.

Sharpe, N.A. & Tepper, J.M. (1998) Postnatal development of excitatory synaptic input to the rat neostriatum: an electron microscopic study. Neuroscience, 84, 1163–1175.

Shepherd, G.M. (2013) Corticostriatal connectivity and its role in disease. Nature reviews, 14, 278–291.

Smith, Y., Galvan, A., Ellender, T.J., Doig, N., Villalba, R.M., Huerta-Ocampo, I., Wichmann, T. & Bolam, J.P. (2014) The thalamostriatal system in normal and diseased states. Frontiers in systems neuroscience, 8, 5.

Smith, Y., Raju, D.V., Pare, J.F. & Sidibe, M. (2004) The thalamostriatal system: a highly specific network of the basal ganglia circuitry. Trends in neurosciences, 27, 520–527.

Sohur, U.S., Padmanabhan, H.K., Kotchetkov, I.S., Menezes, J.R. & Macklis, J.D. (2012) Anatomic and Molecular Development of Corticostriatal Projection Neurons in Mice. Cereb Cortex.

Somogyi, P., Bolam, J.P. & Smith, A.D. (1981) Monosynaptic cortical input and local axon collaterals of identified striatonigral neurons. A light and electron microscopic study using the Golgi-peroxidase transport-degeneration procedure. The Journal of comparative neurology, 195, 567–584.

Taverna, S., Ilijic, E. & Surmeier, D.J. (2008) Recurrent collateral connections of striatal medium spiny neurons are disrupted in models of Parkinson’s disease. J Neurosci, 28, 5504–5512.

Tecuapetla, F., Jin, X., Lima, S.Q. & Costa, R.M. (2016) Complementary Contributions of Striatal Projection Pathways to Action Initiation and Execution. Cell, 166, 703–715.

Tepper, J.M. & Plenz, D. (2006) Microcircuits in the striatum: striatal cell types and their interaction. In: Microcircuits: the interface between neurons and global brain function. MIT Press, Cambridge, Mass.; London.

Tepper, J.M., Sharpe, N.A., Koos, T.Z. & Trent, F. (1998) Postnatal development of the rat neostriatum: electrophysiological, light-and electron-microscopic studies. Developmental neuroscience, 20, 125–145.

Tobach, E., Aronson, L.R. & Shaw, E. (1971) The biopsychology of development. Edited by Ethel Tobach, Lester R. Aronson, and Evelyn Shaw. Academic Press, New York; London.

van der Kooy, D. & Fishell, G. (1987) Neuronal birthdate underlies the development of striatal compartments. Brain research, 401, 155–161.

Venance, L., Glowinski, J. & Giaume, C. (2004) Electrical and chemical transmission between striatal GABAergic output neurones in rat brain slices. The Journal of physiology, 559, 215–230.

Yin, H.H. & Knowlton, B.J. (2006) The role of the basal ganglia in habit formation. Nature reviews, 7, 464–476.

Yu, Y.C., He, S., Chen, S., Fu, Y., Brown, K.N., Yao, X.H., Ma, J., Gao, K.P., Sosinsky, G.E., Huang, K. & Shi, S.H. (2012) Preferential electrical coupling regulates neocortical lineage-dependent microcircuit assembly. Nature, 486, 113–117.

